# Neurodevelopmental origin of seizures in Lowe syndrome

**DOI:** 10.1101/2025.11.28.691107

**Authors:** Sukanta Behera, Pranati Mahajan, Tista Bhattacharya, Padinjat Raghu

**Affiliations:** National Centre for Biological Sciences, GKVK Campus, Bangalore 560065, India; Ashoka University, Rajiv Gandhi Education City, Haryana 131021, India

**Keywords:** Lowe syndrome, OCRL, neurodevelopment, seizures, phosphoinositides, *Drosophila*

## Abstract

Lowe syndrome (LS) is a rare X-linked monogenic disorder resulting from mutations in the *OCRL1* gene that encodes a phosphatidylinositol 4,5-bisphosphate 5’ phosphatase enzyme. Patients with LS exhibit a range of neurological symptoms, including neurodevelopmental delays, hypotonia, febrile seizures, and behavioural abnormalities; however, the cellular and developmental origins of LS remain poorly understood. The *Drosophila* genome encodes a single homolog of OCRL (*docrl*). Here, we report that a germline null allele of *docrl* (*docrl^KO^*) shows heat induced seizures reminiscent of the febrile seizures in LS patients. Cell type specific deletion of *docrl* in neurons was sufficient to recapitulate the heat induced seizures seen in *docrl^KO^* indicating a cell autonomous requirement of *docrl* in neurons to prevent seizures. Temporally controlled deletion of *docrl* showed that heat induced seizure in adults were predetermined by a requirement of *docrl* in neural stem cells during embryonic neurogenesis. Collectively, our findings demonstrate the developmental origin of the neurological manifestations of LS highlighting the need to target potential therapeutic interventions during this developmental time window.

## Introduction

The differential phosphorylation on the inositol head group of phosphatidylinositol can generate seven distinct phosphoinositide species. These seven phosphoinositides provide a unique lipid code to organelle membranes (Balla, 2013) and can serve as signalling molecules in regulating key cellular processes such as receptor signalling, vesicular transport and cytoskeletal function. The cellular levels of these phosphoinositides are set by the activity of specific lipid kinases (enzymes that add a phosphate to the inositol head group) and lipid phosphatases (enzymes that remove a phosphate from the inositol head group). Consequently, mutations in genes encoding these enzymes have been implicated in various diseases including neurodevelopmental disorders, such as Joubert syndrome (*INPP5E*), Parkinson’s disease (*SYNJ*), Charcot-Marie-Tooth disease (*FIG4*) and Lowe syndrome (*OCRL*) [reviewed in (Raghu et al., 2019)].

Lowe syndrome (LS) is a rare X-linked recessive monogenic human disease that affects the eyes, brain and kidneys (Mehta et al., 2014). LS is caused by mutations in the gene *OCRL*; OCRL belongs to the family of 10 inositol polyphosphate 5-phosphatases in vertebrates (Ramos et al., 2019). It encodes a multidomain protein containing four major domains: an N-terminus Pleckstrin homology (PH) domain, a catalytic 5’ phosphatase domain, an ASPM-SPD-2-Hydin (ASH) domain, and a Rho GTPase activating protein (RhoGAP) domain. The OCRL protein is an enzyme that can dephosphorylate both phosphatidylinositol (4,5) bisphosphate [PI(4,5)P_2_] and phosphatidylinositol (3,4,5) trisphosphate [PI(3,4,5)P_3_] with PI(4,5)P_2_ being the preferred substrate (Schmid et al., 2004; Zhang et al., 1995). Consistent with this, previous studies with LS patient derived cells have demonstrated an elevation in the levels of PI(4,5)P_2_ (Akhtar et al., 2022; Sharma et al., 2024; Wenk et al., 2003). In humans, the *OCRL* gene encodes two isoforms: isoform A and isoform B. The two isoforms are expressed across all tissues, except the brain. The brain expresses only isoform A, which has 8 additional amino acids that improve its clathrin binding(Choudhury et al., 2009).

Clinically, the eye manifestation in LS patients is typically a congenital cataract and in the case of the kidney, a proximal tubule reabsorption defect. The brain manifestations of LS can be many and diverse. Reported manifestations include hypotonia, joint hypermobility and areflexia suggesting severe psychomotor defect, delayed development, speech defects and intellectual disability (Loi, 2006; Röschinger et al., 2000; Şimşek et al., 2011). LS patients can also manifest behavioural defects such as stubbornness, temper tantrums, aggressive and repetitive behaviour (Arron et al., 2011; Kenworthy et al., 1993). In addition, LS patients are prone to febrile seizures that resemble Lennox-Gastaut type seizures (Giannakopoulos et al., 1990). Brain imaging studies have shown the presence of increased periventricular density (O’Tuama and Laster, 1987), cystic lesions (Erdoǧan et al., 2007), white matter abnormalities and increased myo-inositol peaks suggesting gliosis and demyelination (Sener, 2004; Yuksel et al., 2009). Neuropathological studies have shown cerebral atrophy (Giannakopoulos et al., 1990), and slow nerve conduction (Kornfeld, 1975).

Although many studies have sought to address the renal phenotypes of LS, there have been limited animal models that have enhanced our understanding of the brain phenotype of these patients. A notable study is of *OCRL* depletion in a zebrafish model; Ramirez et.al studied mutants in zebrafish *OCRL* (Ramirez et al., 2012) and reported increased susceptibility to heat-induced seizures, reduced brain size, increased apoptosis and features suggestive of gliosis (Ramirez et al., 2012). In contrast, mouse models for *OCRL*, have failed to manifest brain phenotypes (Festa et al., 2019) likely due to a functional overlap between *OCRL* and its paralog *INPP5B* (Jänne et al., 1998). Most recently, Sharma et.al (Sharma et al., 2024) have reported altered physiological functions and brain development in a LS patient iPSC derived brain organoid model; this study reported delayed development of neuronal activity, altered levels of GFAP, and Notch signalling that were dependent on the levels of PI(4,5)P_2_ in neural cultures.

The *Drosophila* genome contains a single gene for OCRL (*docrl)*(Balakrishnan et al., 2015). Previous studies have reported phenotypic effects of *docrl* depletion. Del Signore et.al reported effects of *docrl* mutants on haemocyte cell biology and proposed that this might affect the brain in a non-cell autonomous manner although no brain phenotypes were reported in this study. Ramesh et.al also analysed a CRISPR engineered deletion, *docrl^KO^* (Trivedi et al., 2020) and reported phenotypes related to nephrocyte function (Ramesh et al., 2024); however, no brain phenotype was reported in this study.

In this study, we report that loss of *docrl* leads to heat induced seizure activity in adult flies. Cell type specific deletion revealed that this phenotype arises from a requirement in neurons but not glia. Stage specific deletion experiments established that *docrl* is required in immature neurons and neuroblasts but not in mature neurons. Seizure activity in adult flies arises from depletion of *docrl* in the 1^st^ phase of neurogenesis, suggesting a role in early neurodevelopment. Lastly, at a physiological level, *docrl* depleted animals show synchronous firing of motor neurons similar to that seen in other seizure sensitive mutants like *para^bss1^*. Thus, our study establishes the requirement of *docrl* in early *Drosophila* neurodevelopment to prevent heat induced seizures in adults.

## Results

### *docrl* depletion increases susceptibility to heat-induced seizures in adult flies

We have previously noted that the *docrl* germline knockout (*docrl^KO^*) (Trivedi et al., 2020) flies are pupal lethal (Ramesh et al., 2024). Culturing these flies in jaggery enriched medium led to enhanced larval growth, contributing to some flies escaping pupal lethality and emerging as escaper male adult flies. These adult flies are smaller in size and weaker than the controls. We confirmed the presence of CRISPR deletion through detection of a PCR band of 555bp in the *docrl^KO^* male flies; indicating the presence of *docrl^KO^* genomic deletion **[Fig1 E]**.Western blotting confirmed that these males are dOCRL protein null **[Fig1 F]**. The availability of such hemizygous adult escaper *docrl^KO^* males allowed us to study adult behaviour in *Drosophila*.

LS patients show neurological phenotypes such as psychomotor defects, behavioural and cognitive defects seizures, delays in development (Bökenkamp and Ludwig, 2016; Mehta et al., 2014). Of these, seizure behaviour is seen in ca. 50% of LS patients (Loi, 2006). These vary in various types from infantile febrile (commonly seen), tonic-clonic, myoclonic, and atonic seizures (Erdoǧan et al., 2007).

Using these germline *docrl^KO^* adult flies, we performed heat-induced seizure behaviour assay at 42^°^C [**Fig 1A**]. Briefly, adult flies in empty glass vials are subjected to a temperature of 42^°^C; under these conditions, only a small proportion of adults fall to the bottom of the vial (referred to as falling frequency) [**Fig 1B**]. However, under the same conditions, *docrl^KO^* flies showed a statistically significant increase in falling frequency [**Fig 1B**]. In addition, while only a few control flies showed post-seizure paralysis, ca. 78% *docrl^KO^*flies showed paralysis **[Fig1 C]**. When flies were returned to room temperature after the heat shock, *docrl^KO^* animals recovered from paralysis with a t_1/2_ of ca. 50s [**Fig 1D**]. These phenotypes of *docrl^KO^* could be rescued by introduction of an X-chromosome duplication encompassing the *docrl* locus [**Fig 1 B-D].** A representative video of the assay is shown in **[Suppl Video 1]**. These findings suggest that loss of *docrl* leads to heat induced seizures.

**Figure 1:**
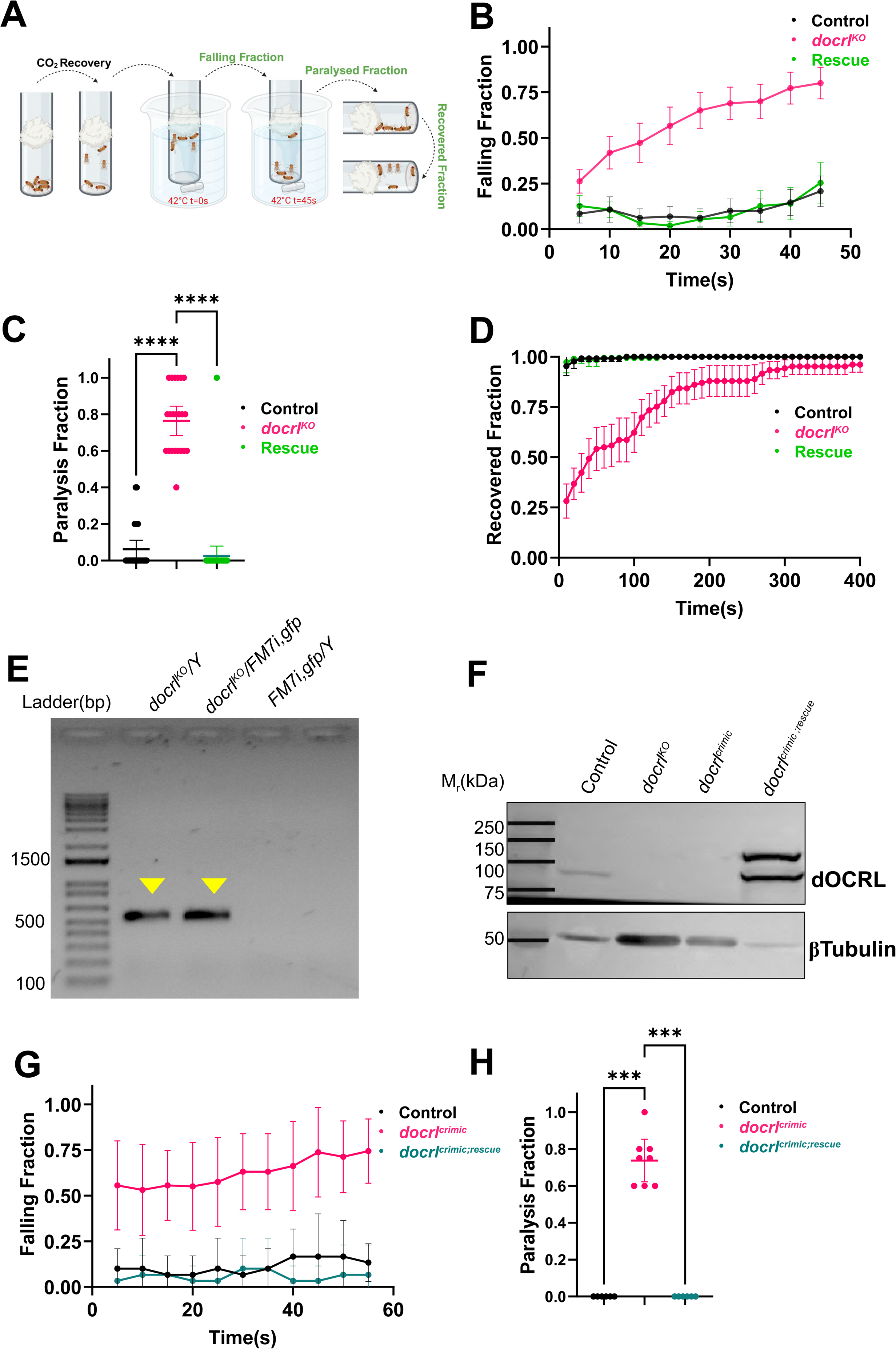
Heat induced seizure assays in *docrl^KO^*. **(A)** Schematic representation of heat induced seizure assay. Empty glass vials with flies from the point of collection to the end of the assay are shown. The sequence of steps is indicated by dotted arrows. The temperature of the water bath is indicated. The parameters measured in this behavioural assay are marked in green (Created with BioRender.com). Behavioural parameters calculated for various genotypes of 2-4 day old adult flies. The black colour represents control (*y^1^v^1^/Y) (N=13 sets (each set has 5 flies), n=65 flies),* the pink colour represents (*docrl^KO^ /Y) (N=11 sets, n=55 flies),* and the green colour represents the rescue files (*docrl^KO^/Y;;Dp(1:3)DC402) (N=19 sets, n=95 flies).* **(B)**The X axis represents the time in seconds; the Y axis represents the falling fraction. Error bars represent mean falling frequency per trial ±95% CI (Confidence Intervals). **(C)** Fraction of flies paralysed at the end of the heat shock period. Y axis represents the paralysis fraction. Each point is showing fraction of flies paralysed per trial. The graph shows the mean paralysis fraction per trial±95% CI. Statistics used : Kruskal-Wallis test post hoc Dunn’s multiple correction,**** represents p value <0.0001. **(D)** Fraction of flies recovered from paralysis as a function of time. X-axis represents time in seconds. Y axis represents the recovered fraction. Error bars represent mean recovered fraction per trial ±95% CI **(E).** Representative PCR analysis to detect the *docrl^KO^* deletion from DNA extracts of adult flies. Lane 1 - hemizygous *docrl^KO^ /Y* adult; lane 2 - heterozygous balanced *docrl^KO^/FM7i,gfp* and lane 3 - negative control *FM7i,gfp/Y* flies. A diagnostic 555 bp band for the *docrl^KO^* deletion is shown by the yellow arrowhead. **(F)** Western blot of dOCRL protein levels in *docrl* mutants. Tubulin is used as a loading control. The genotypes analysed are: control (*y^1^v^1^/Y), docrl^KO^* (*docrl^KO^ /Y), docrl^crimic^ (docrl^crimic^/Y)* and *docrl^crimic;rescue^ (docrl^crimic^; UAS-docrl::gfp).* **(G,H)** Heat induced seizure quantified in 2-4 day old adult flies of *docrl^crimic^*. The black colour represents internal control which expresses endogenous dOCRL (*docrl^crimic^/Y -P3 RFP*) (N=3 sets,n=15 flies), the pink colour represents (*docrl^crimic^ /Y*) (N=4 sets, n=20 flies), and the dark green colour represents the rescue files *docrl^crimic;rescue^*(*docrl^crimic^/Y;UAS-docrl::gfp)* (N=3 sets, n=15 flies). In G, the X axis represents the time in seconds; Y axis represents the falling fraction. Error bars represent mean falling frequency per trial ±95% CI. In H the fraction of flies paralysed at the end of the heat shock period is shown. Y axis represents the paralysis fraction. Each point is the fraction of flies paralysed per trial. The graph shows the mean paralysis fraction per trial±95% CI. Statistics used : Kruskal-Wallis test post hoc Dunn’s multiple correction, **** represents p value <0.0001.

In addition, we used the *docrl^crimic^* allele, which generates an early truncation of the *docrl* mRNA and expresses GAL4 from the endogenous *docrl* promoter (Lee et al., 2018). These flies exhibit larval phenotypes similar to the *docrl^KO^* allele and on Western blot analysis, we found that they are protein-null [**Fig 1F**]. When grown on jaggery enriched medium, we obtained a few adult male escapers of *docrl^crimic^*. The flies exhibited the heat induced seizures seen in *docrl^KO^* [**Fig 1 G,H]**. Expression of wild type *docrl* in *docrl^crimic^* using *UAS-docrl::gfp* (referred as *docrl^crimic;rescue^*) [**Fig 1F**] rescued the larval lethality, and the resulting adult male flies did not exhibit increased seizure susceptibility [**Fig 1 G,H]**.

### *docrl* depletion in neurons is sufficient to trigger heat induced seizures

Seizures occur as a result of neuronal hyperexcitability that may arise as a consequence of brain intrinsic factors or secondary to systemic factors such as metabolic dysregulation or altered immune cell function (Shen et al., 2016). Within the brain itself, primary defects in neurons can result in seizures; alternatively changes in glial cell function may also lead to neuronal hyperexcitability, manifest as seizures (Kunduri et al., 2018; Pitkänen et al., 2017; Wang et al., 2004). To understand the cellular origin of heat induced seizures in *docrl^KO^* we performed a screen in which *docrl* was selectively depleted in various tissues while testing for seizures in adult flies. Tissue specific knockout of *docrl^KO^* was achieved using a dual guide RNAs (dgRNAs) for *docrl* and individual GAL4 lines driving UAS-Cas9 expression thus, generating tissue specific *docrl* knockout [**Fig 2 A]**(Trivedi et al., 2020).

**Figure 2:**
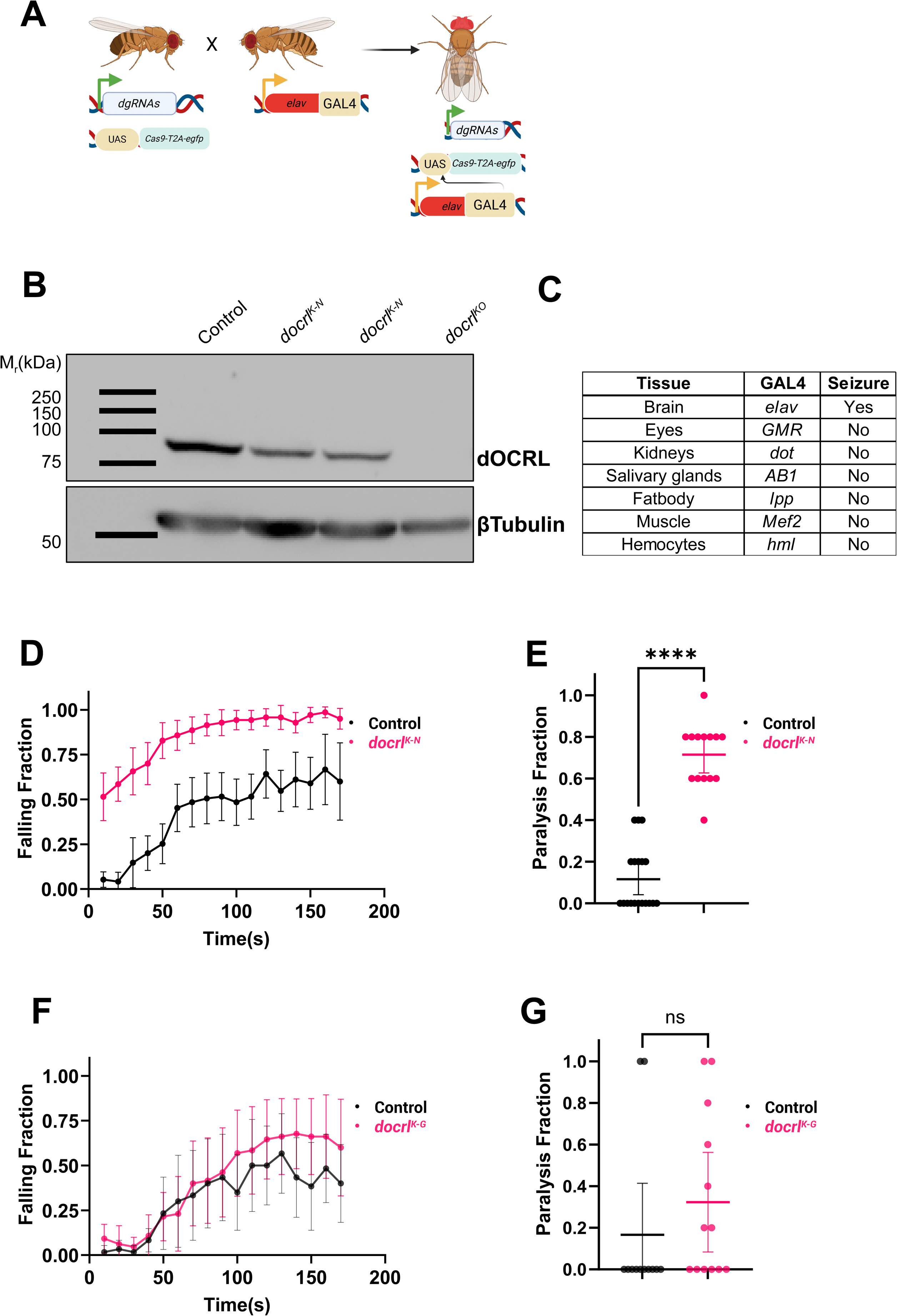
Heat induced seizure assays in tissue specific *docrl* deletion. **(A)** Schematic representation of tissue specific *docrl* knockout using CRISPR/Cas9 genome engineering. Flies expressing the dual guideRNA for *docrl* (*dgRNAs*) ubiquitously and a *UAS-Cas9-T2A-egfp* transgene are shown. Flies expressing a specific GAL4 (*elav-GAL4)* are shown, the progeny with genotype is indicated (Created with BioRender.com). **(B)** Western blot assessing the depletion of *docrl* protein in brain tissue extracts of tissue specific deletion flies. The genotypes analysed are: control (*elav-GAL4;dgRNAs),* neuron specific knockout *docrl^K-N^(elav-GAL4;dgRNAs;UAS-Cas9-T2A-egfp).* Germline *docrl^KO^* flies are used as a control for the specificity of the dOCRL band detected. Tubulin levels are used as a loading control. **(C)** Table showing various tissue screened for seizure susceptibility using tissue specific CRISPR/Cas9 deletion of *docrl*. Detection of seizure behaviour is indicated by yes, no-indicates no seizures. The identity of the GAL4 lines used for each tissue is indicated. **(D,E)**Heat induced seizure assay with 2-4 days old adult flies carrying docrl knockout in neurons (*docrl^K-N^)*. The genotypes of control (*elav-GAL4;dgRNAs*)*(N=10 sets, n=50 flies)* and *docrl^K-N^(elav-GAL4;dgRNAs;UAS-Cas9-T2A-egfp)* (N=7 sets, n=35 flies). **(D)** The X axis represents the time in seconds; Y axis represents the falling fraction. Error bars represent mean falling fraction±95% CI, **(E)** Y-axis indicates the paralysed fraction. Each point represents the paralysis fraction per trial. Error bars represents mean paralysis fraction±95% CI, Statistics used: Mann-Whitney U test, **** represents p value <0.0001. (**F, G)** Heat induced seizure assay with 2-4 days old adult flies having *docrl* knockout in glia. The genotypes are control (*dgRNAs;repo-GAL4*) (N=6 sets, n=30 flies) and *docrl^K-G^(dgRNAs;UAS-Cas9-T2A-egfp/repo-GAL4)* (N=7 sets, n=35 flies). The X axis represents the time in seconds; Y axis represents the falling fraction in F and paralysis fraction in G respectively. The graph shows mean±95% CI, Statistics used : Mann-Whitney U test, ns represents p value >0.05.

To validate the effectiveness of our knockout system, we expressed Cas9 using the ubiquitously expressed *da (daughterless)-GAL4 (docrl^K-D)^.* PCR analysis of fly body extracts of *docrl^K-D^*showed a band demonstrating the deletion of a part of the *docrl* gene using these dgRNAs **[Suppl Fig 1 A]**. Western blot analysis of *docrl^K-D^*extracts showed a reduction in dOCRL protein levels **[Suppl Fig 1B].** In behaviour assays, *docrl^K-D^* flies recapitulate the heat induced seizure susceptibility seen in *docrl^KO^ flies* **[Suppl Fig 1 C-E]**.

We knocked out *docrl* in neurons using the pan neuronal driver, *elav-GAL4 (docrl^K-N^)* and glia using the pan glial driver *repo-GAL4 (docrl^K-G^)* and validated the depletion in dOCRL protein levels in adult fly heads in *docrl^K-N^* by Western blotting [**Fig 2 B]**. Behaviour assays showed enhanced heat induced seizure [**Fig 2 D]** with ca. 75% paralysis [**Fig 2 E]** in *docrl^K-N^* but there was no significant change in *docrl^K-G^*[**Fig 2 F,G]**. A representative video of heat induced seizure assays with pan neuronal and pan glial *docrl* knockout are shown in **[Suppl Video 2-3]**.

### *docrl* function in non-brain tissues does not contribute to heat induced seizures

To test the possible role of a non-cell autonomous origin of *docrl* depletion on seizure susceptibility, we depleted *docrl* in the eye and kidney, that are affected in LS patients (Mehta et al., 2014). Depletion in *Drosophila* nephrocytes (*docrl^K-R^)*, which are analogous to the vertebrate kidney, and *Drosophila* eyes (*docrl^K-E^)* did not show any significant change in the falling frequency or paralysis frequency compared to controls **[Supp Fig 2-A,B]**.

We also tested the potential role of other organs like salivary glands (*docrl^K-SG^)* and fat body (*docrl^K-FB^)*, which are major secretory organs implicated in communication to the brain (Li et al., 2022; Rajan and Perrimon, 2012) and found no impact of *docrl* depletion in these tissues on falling frequency or paralysis **[Suppl Fig 2 C-F]**. A previous study had suggested that loss of *docrl* in haemocytes might lead to phenotypes in the nervous system (Del Signore et al., 2017)To test this, we selectively depleted *docrl* in haemocytes (*docrl^K-H^)* **[Suppl Fig 2 D]** and found no seizure phenotype.

The peripheral nervous system (PNS) is important in sensing noxious stimuli like heat, light, and electricity via multidendritic sensory neurons (Kilo et al., 2021). We employed *sens-GAL4* which expresses in a majority of the PNS (Nolo et al., 2000) and found no evidence of seizure susceptibility (data not shown). A summary of all tissues screened for heat induced seizures is shown in [**Fig 2 C]**. Together, these data suggest that loss of *docrl* in the neurons of the central nervous system contributes to heat induced seizures.

### *docrl* depletion in mature neurons is insufficient to generate heat induced seizures

The *Drosophila* central nervous system is composed of multiple types of neurons and *docrl* is expressed in most of them. To test if the *docrl* function is specifically required in any particular neuronal subtype in respect to susceptibility to heat-induced seizures, we depleted the gene in each subtype. Selective depletion of *docrl* individually in cholinergic, glutamatergic, dopaminergic, serotonergic, GABAergic, octopaminergic, and peptidergic neurons did not result in a significant increase in falling frequency in the heat induced seizure assay **[Suppl Fig 3 A-F].** This suggests that loss of *docrl* in one neuronal subtype is insufficient to induce heat susceptibility.

Considering that these neuronal subtypes specific GAL4 are expressed in mature neurons. We used another pan neuronal driver *nSyb (neuronal Synaptobrevin)* that is expressed in most of the mature neurons (Marques et al., 2023) to deplete *docrl (docrl^K-nSyb^)*. The *docrl^K-nSyb^* flies did not show an increased susceptibility to heat induced seizures in 2-4 days old flies [**Fig 3 B]**. We also aged the flies to 11-13 days [**Fig 3 C]** to remove any effect of perdurance of dOCRL protein but still we saw no significant difference in seizure susceptibility. Therefore, depletion of *docrl* in mature neurons is insufficient to confer susceptibility to heat induced seizures.

**Figure 3:**
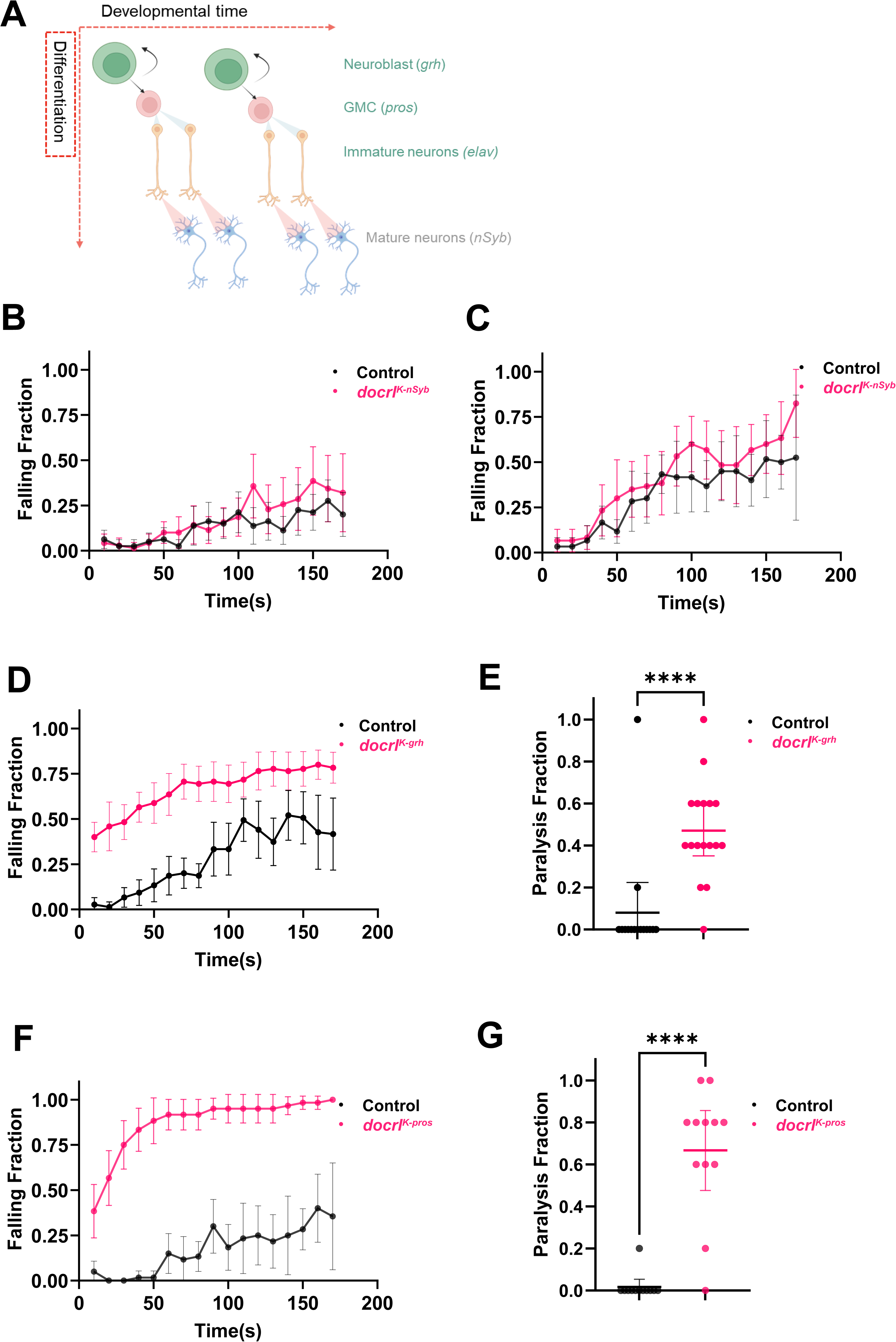
**(A)** Representation of *Drosophila* neuroblast division across the differentiation axis. The neuroblast (NB) divides asymmetrically to form another neuroblast and a ganglion mother cell (GMC). These GMCs then undergo a terminal division to generate immature neurons. The immature neurons then mature by forming functional synapses and expressing neurotransmitters specific to their final neuronal identity (Created with BioRender.com). Transcripts unique for each cell type are shown NB (*grh*); GMC (*pros*); immature neuron (*elav)*; mature neuron (*nSyb*) are marked in round brackets. Results of heat induced seizure assays in mature neurons **(B)**1-4 days old flies and **(C)** 11-13 days old flies. The genotypes of control (*dgRNAs;nsyb-GAL4*) (N=6-8 sets, n=30-40 flies) and *docrl^K-nSyb^(dgRNAs;grh-GAL4/UAS-Cas9-T2A-egfp)* (N=6-7 sets, n=30-35 flies). **(D,E)** Results of heat induced seizure assays in 2-4 days old adult flies upon *docrl* knockout in neuroblasts. The genotypes are control (*dgRNAs/grh-GAL4*) (N=8 sets, n=40 flies) and *docrl^K-grh^ (dgRNAs/grh-GAL4;UAS-Cas9-T2A-egfp)* (N=9 sets, n=45 flies). **(F,G)** Results of heat induced seizure assays in 2-4 days old adult flies upon *docrl* knockout in GMC. The genotypes are control (*dgRNAs;pros-GAL4*) (N=6 sets, n=30 flies) and *docrl^K-pros^ (dgRNAs;UAS-Cas9-T2A-egfp/pros-GAL4)* (N=6 sets, n=30 flies). In B,C,D and F, the X axis represents the time in seconds, and the Y axis represents the falling fraction. Error bars represent mean±95%CI. In E and G Y-axis represents the paralysis fraction. The bar represents mean±95%CI, each point represents the paralysis fraction per trial. Statistics used: Mann-Whitney U test, **** represents p value <0.0001

### *docrl* depletion in neuronal precursors is required for seizure susceptibility

While depletion of docrl in mature neurons (*docrl^K-nSyb^)* did not generate heat induced seizures, the use of e*lav-GAL4* that expresses in immature, post-mitotic neurons was able to do so *(docrl^K-N^)*. This suggests that heat induced seizures in *docrl^KO^* arise from a requirement of dOCRL function during neuronal development. In *Drosophila*, neurons are born from a neuroblast (NB) that divides to form a ganglion mother cell (GMC) which further generates immature neurons [**Fig 3 A]** (Homem and Knoblich, 2012). We tested if loss of *docrl* in neuroblast or ganglion mother cells would result in heat induced seizures in adult flies. Remarkably, we found that depletion of *docrl* in neuroblasts *(docrl^K-grh^)*, marked by *grainyhead (grh),* the last transcription factor expressed in neuroblasts(Almeida and Bray, 2005; Isshiki et al., 2001) of *Drosophila* neurogenesis led to heat induced seizures [**Fig 3 D, E]**. Likewise, depletion in ganglion mother cells *(docrl^K-pros^)*, marked by *prospero* (Crews, 2019), also resulted in heat induced seizures [**Fig 3 F, G]**, recapitulating phenotypes seen with *docrl^K-N^*. These results suggest that *docrl* depletion during neuronal development is necessary to endow susceptibility to heat induced seizures.

### *docrl* is required during the early stages of neurodevelopment

During *Drosophila* development, neurogenesis occurs in two distinct phases, a 1^st^ phase during embryogenesis to 1^st^ instar larvae and a 2^nd^ phase from 1^st^ instar to pupariation following which neuronal remodelling occurs during pupal development [**Fig 4 A]** (Homem and Knoblich, 2012).

**Figure 4:**
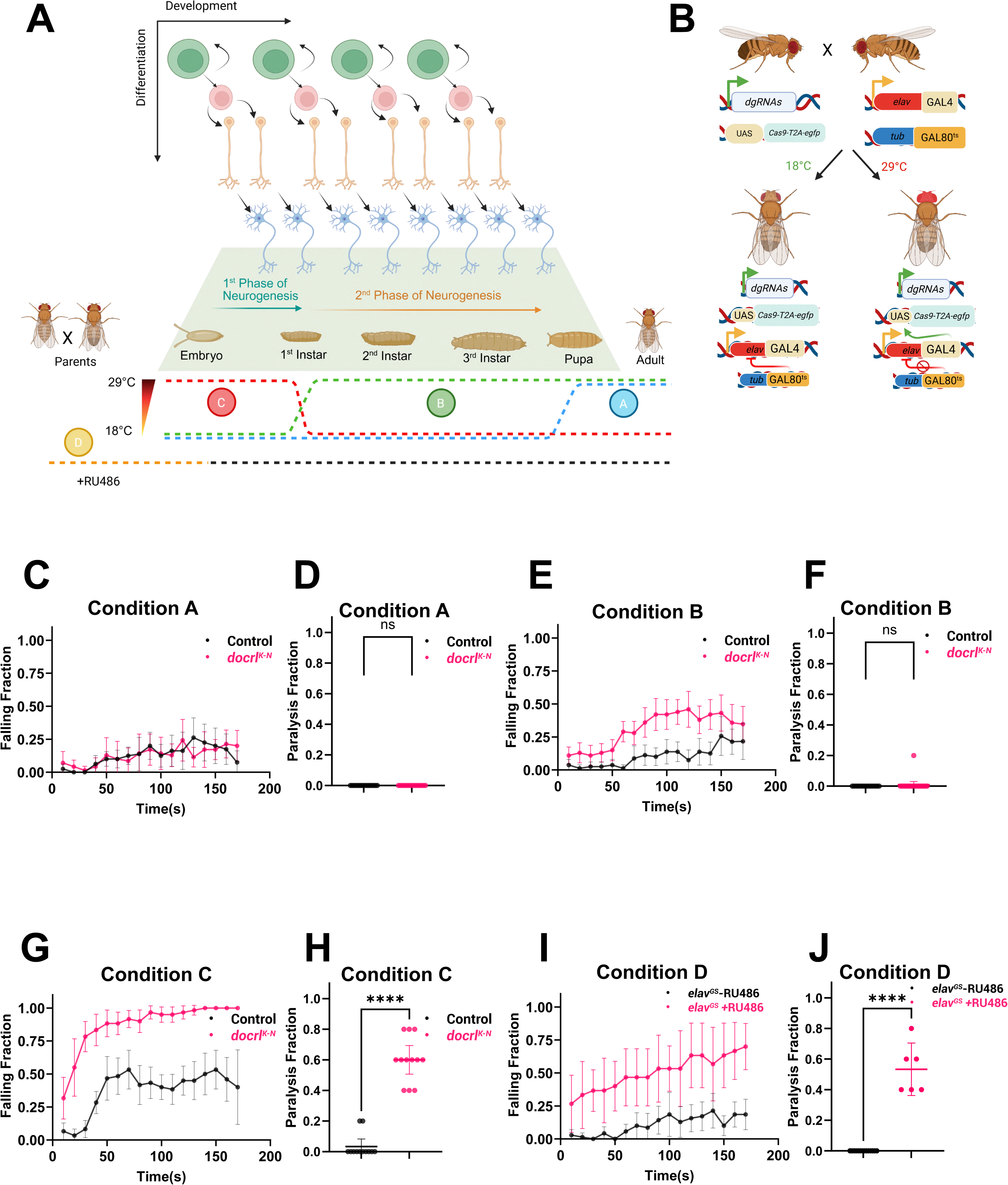
Temporal requirement of *docrl* during development. **(A)** Pictorial representation of the phases of *Drosophila* neurodevelopment. Neurogenesis occurs in two distinct phases: the first phase takes place during the embryonic and 1^st^ instar larval stages (marked by the blue arrow), and the second phase occurs from the late first instar to the pupal stage (marked by the orange arrow). During pupal metamorphosis, a period of significant neuronal remodeling occurs to form the adult brain. At the bottom, experimental condition for *docrl* knockout across various developmental stages using the TARGET approach is shown. The temperature is elevated from 18^°^C to 29^°^C to inactivate the GAL80^ts^. Depletion of *docrl* during 1^st^ phase of neurogenesis (embryo to 1^st^ instar) shown in red dotted lines and marked as condition C. Depletion of *docrl* during 2^nd^ phase of neurogenesis (late 1^st^ instar to pupa) shown in green dotted lines and marked as condition B. Depletion of *docrl* from pupal stage onwards is shown in blue dotted lines and marked as condition A. Condition D shows an alternative approach utilizing the Geneswitch GAL4 system. To achieve *docrl* depletion during the embryonic stage, the parental cross is fed with either 15μM RU486 or ethanol (vehicle control) (as indicated by the orange dotted lines) (Created with BioRender.com). **(B)** Schematic of the system used to achieve temporal gene knockout by combining the GAL4/GAL80^ts^ system with CRISPR/Cas9. At a permissive temperature of 18^°^C (shown in green), the GAL80^ts^ repressor is active, binding to GAL4 and blocking its activity. This prevents the expression of the Cas9 nuclease. When the temperature is shifted to a restrictive temperature of 29^°^C(shown in red), GAL80^ts^ becomes inactive, allowing GAL4 to drive the expression of Cas9 in the desired tissue (Created with BioRender.com). **(C-D)** Heat induced seizure assay in 2-4 days old adult flies with *docrl* depletion induced from the pupal stage onwards (condition A marked in A). **(E-F)** Heat induced seizure assay in 2-4 days old adult flies with *docrl* knockout from late 1^st^ instar to pupal stage (corresponding to 2^nd^ phase of neurogenesis condition B marked in A). **(G-H)** Heat induced seizure assay in 2-4 days old adult flies with *docrl* knockout from embryonic to 1^st^ instar larvae (corresponding to 1^st^ phase of neurogenesis condition C marked in A). The genotypes of the controls: *elav-GAL4; dgRNAs;tub-GAL80^ts^ (*N=6-8 sets, n=30-40 flies) and the knockouts are *elav-GAL4; dgRNAs;UAS-Cas9-T2A-egfp/tub-GAL80^ts^(*N=6-10 sets, n=3-50 flies). **(I-J)** Heat induced seizure assay in 2-4 days old adult flies with *docrl* knockout from embryonic stage to 1^st^ instar is induced by Geneswitch GAL4 system marked as condition D in A. The genotypes of the ethanol fed controls *elav^GS^-RU486* are *elav^GS^;dgRNAs;UAS-Cas9-T2A-egfp (*N=7 sets, n=35 flies) and the 15µM RU486 fed knockouts *elav^GS^+RU486* are *elav^GS^;dgRNAs;UAS-Cas9-T2A-egfp (*N=3 sets, n=15 flies). In C,E,G,I, the X axis represents the time in seconds and the Y axis represents the falling fraction. The graph represents the mean falling fraction ±95%CI. In D,F,H,J, the X axis represent the genotype, and the Y axis represents the paralysis fraction. The bar represents mean paralysis fraction ±95%CI, Statistics used: Mann-Whitney U test, **** represents p value <0.0001 and ns represents p value >0.05.

We investigated the requirement of *docrl* during these phases of neurogenesis. To do this we used cell type expression of Cas9 using specific GAL4 lines but controlled the transcriptional activity of GAL4 using the temperature sensitive repressor GAL80^ts^ [**Fig 4B**][TARGET approach (McGuire et al., 2004)]. We initially activated GAL4 expression in neurons during pupal development, thereby depleting *docrl* during this period. Behavioural assays did not show a significant increase in falling frequency or paralysis [**Fig 4 C-D]**.

When *docrl* was depleted during the 2^nd^ phase of neurogenesis, i.e., during 1^st^ instar to pupal development, we noted a modest increase in susceptibility to heat induced seizure but no paralysis [**Fig 4 E-F]**. However, depletion of *docrl* during 1^st^ phase of neurogenesis showed a significant increase in falling frequency and paralysis compared to control [**Fig 4 G-H]** thus, recapitulating the defect seen in *docrl^K-N^* [**Fig 2 D-E]**. In an alternative approach we temporally controlled GAL4 expression using a drug inducible Geneswitch GAL4 (Osterwalder et al., 2001), *elav^GS^.* Feeding 15mM of RU486 (a progesterone hormone analog) to adult parents led to *docrl* depletion during embryogenesis and resulted in increased susceptibility to heat induced seizures [**Fig 4 I-J]**. These findings suggest that dOCRL function during the first phase of neurogenesis underlies the susceptibility to heat induced seizures in adult flies.

### dOCRL protein is highly expressed in developing neural tissues

We investigated the expression pattern of dOCRL across *Drosophila* neurodevelopment. For this UAS-*docrl::gfp* was expressed using *docrl^crimic^* (Lee et al., 2018) where dOCRL::GFP is expressed under the endogenous *docrl* promoter. We noted the expression of dOCRL::GFP in the ventral nerve cord and other peripheral tissues of late embryonic stages [**Fig 5A**]. In the wandering 3^rd^ instar larval brain [**Fig 5B**], we noted higher expression of dOCRL::GFP in developing optic lobes, neurons, and progenitor cells (marked by absence of ELAV protein). In the adult brain higher expression of dOCRL::GFP was noted in anatomical locations such as the optic and antenna lobes (Nichols, 2006) compared to other adult brain regions [**Fig 5C**]. These regions start to develop during the late 3^rd^ instar and pupal stages during metamorphosis (Jefferis et al., 2002; Özel et al., 2021).Thus, dOCRL is expressed ubiquitously in all brain cells but has higher expression in the developing brain regions.

**Figure 5:**
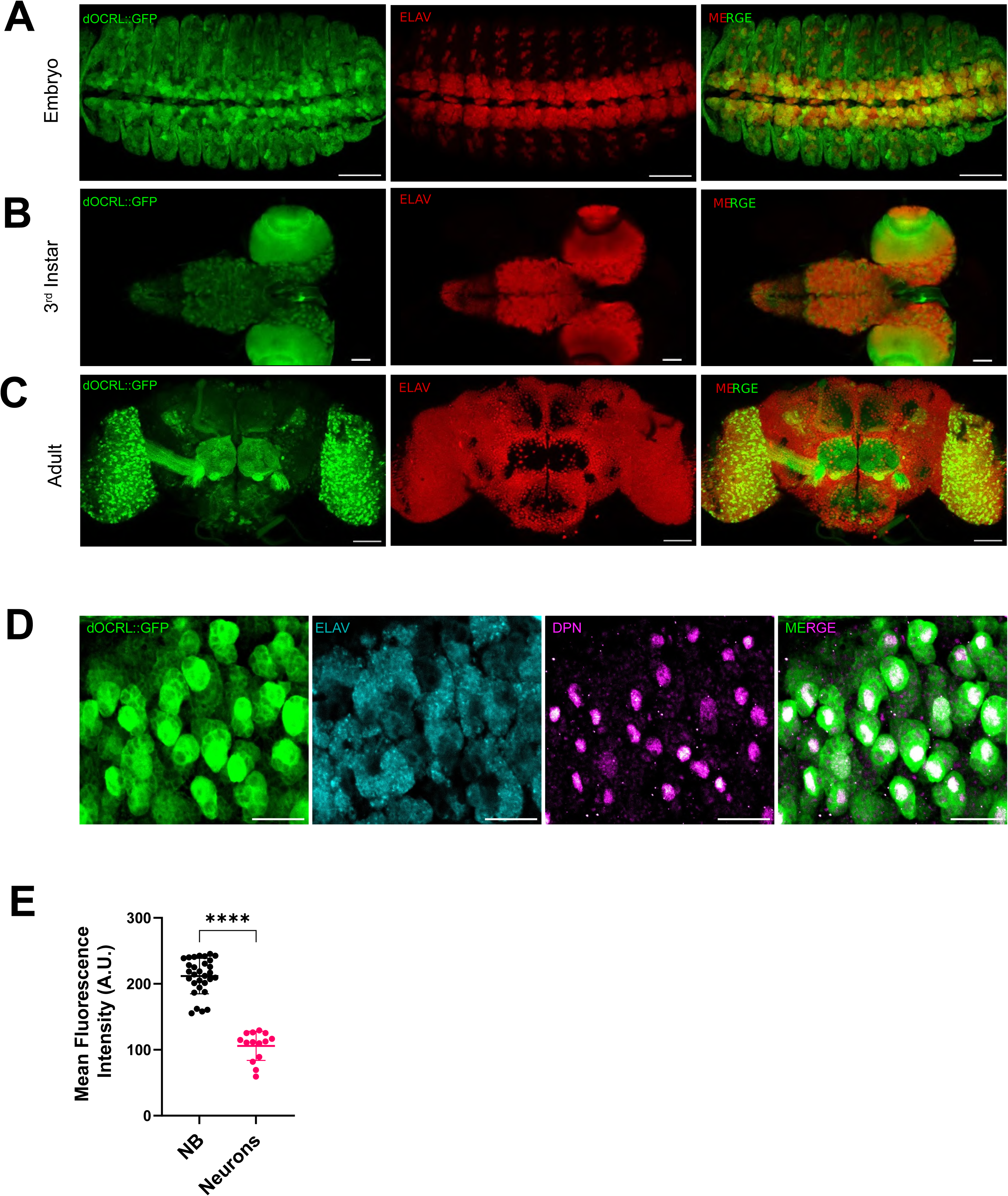
Localisation of *docrl* during development. Confocal images show the expression of dOCRL in the *Drosophila* brain across **(A)** embryo **(B)** 3^rd^ instar larvae and **(C)** adult. The green channel represents dOCRL::GFP protein levels expressed using *docrl^crimic,^* the red channel represents the post mitotic neurons marked by Elav. **(A-C)** Scale bar: 50µm **(D)** A confocal image showing expression of dOCRL in the 3^rd^ instar larval ventral nerve cord (VNC). The green channel represents dOCRL::GFP expression using *docrl^crimic^,* the cyan channel shows the post mitotic neurons marked by Elav, the magenta channel shows the neuroblasts marked by Deadpan. Scale bar: 20µm. **(E)** Quantification of dOCRL::GFP levels in the neuroblast and the neurons from the image in D. The X axis represents the cell type and the Y axis represents the mean fluorescence intensity in arbitrary units (A.U.). Each point is an ROI used to calculate the levels of dOCRL::GFP for each cell type. Statistics used : Mann-Whitney U test, (N=1 brain n=14-30 ROIs), **** represents p value <0.0001.

To confirm our observations we stained a 3^rd^ instar larval brain expressing dOCRL::GFP under *docrl^crimic^* with Deadpan (antibody that marks the neuroblast) and Elav (antibody that marks the post-mitotic neurons) antibodies [**Fig 5 D]** and quantified the dOCRL::GFP intensities of neuroblast and neurons. We saw significantly higher dOCRL levels in neuroblast compared to neurons [**Fig 5 E]**.

### Loss of *docrl* increases synchrony in neuronal activity

We sought to determine a physiological correlate of the seizure activity seen in *docrl^KO^*. Since our data indicates that *docrl* function during embryonic development leads to the seizure activity seen in adult flies, we investigated the physiological state of 3^rd^ instar larval brain neurons using intracellular calcium imaging. It is well established that in wild-type larvae, the fictitious locomotor activity of two adjacent ipsilateral segments in the isolated 3^rd^ instar larval brain has a lag of 800-1000ms whereas seizure sensitive mutants of *the para* gene like *para^bss1^* and *para^GEFS+^* show smaller lags and fire together leading to synchrony. (Streit et al., 2016). To measure activity through changes in intracellular calcium levels, we expressed GCaMP5G, a genetically encoded calcium indicator in the larval motor neurons of the *docrl^KO^* and control larvae. We measured the fictitious calcium waves in the isolated 3^rd^ instar larval brain [**Fig 6 A,C,D]** and used cross correlation analysis to establish whether two ipsilateral neuronal pairs are synchronous or asynchronous [**Fig 6 B]**.

**Figure 6:**
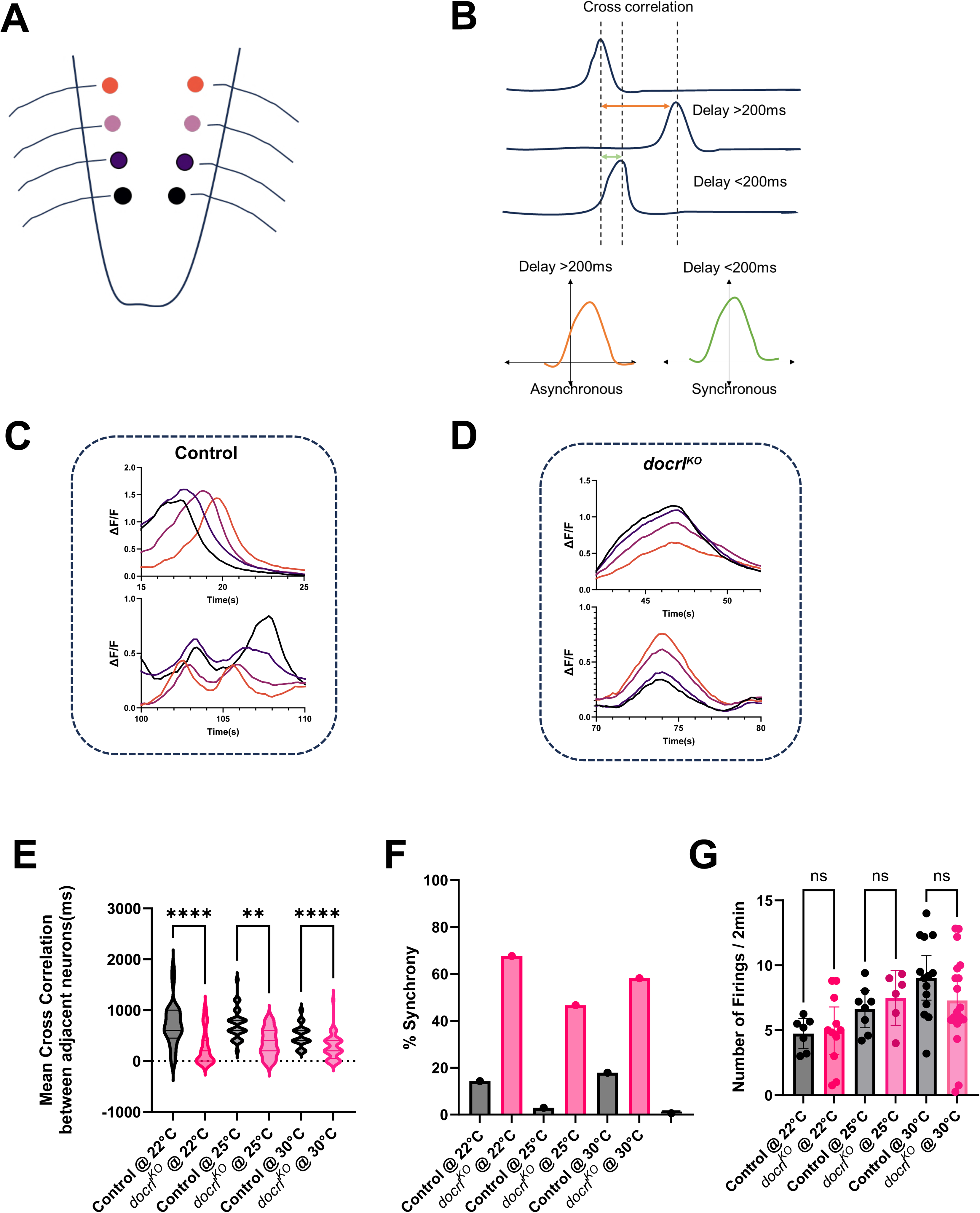
Calcium signalling physiology *docrl^KO^*. Measurement of neuronal activity by intracellular calcium imaging. **(A)** A schematic representation of a *Drosophila* third-instar larval brain (ventral nerve cord). The colour-coded circles represent the regions of interest (ROIs) selected for analysis. Representative calcium traces from individual VNC cells of **(C)** control (*y^1^v^1^; ok6-GAL4; UAS-GCaMP5G*) and **(D)** *docrl^KO^* (*docrl^KO^; ok6-GAL4; UAS-GCaMP5G*). The X-axis represents time in seconds, and the Y-axis represents the normalised change in fluorescence (ΔF/F), calculated relative to the minimum fluorescence of the total frames. **(B)** A representation of the cross-correlation analysis performed between two adjacent ipsilateral neurons. A cross-correlation value greater than 200 milliseconds (ms) indicates asynchronous activity, while a value less than 200 ms is categorized as synchronous. **(E)** Absolute quantification of the mean cross correlation across various ipsilateral neuron pairs. The black represents the control, and pink represents the *docrl^KO^* (genotypes are described above). The X axis represents temperature groups and the Y axis represents the mean cross-correlation coefficient in ms. N=6-10 larval brains per condition, statistics used: Kruskal Wallis test post hoc Dunn’s multiple corrections, **** represents p value <0.0001 and ** represents p value < 0.002. **(F)** A representation of the percentage of neurons showing synchronous vs non-synchronous behaviour; genotypes and temperatures are shown. Fractions calculated from data shown in E. The X-axis represents the temperatures studied and the Y-axis represents the percentage of synchronous events over all recorded events. **(G)** Quantification of the number of calcium peaks in control and *docrl^KO^ at* three different temperatures. The X axis represents the temperature groups and genotypes; the Y axis represents the total number of firing events per 2 minutes of activity recorded. N=6-10 larval brains per condition, statistics used: Kruskal-Wallis test post hoc Dunn’s multiple corrections, ns represents p value >0.1234.

We observed that in contrast to control flies, intracellular calcium waves of adjacent segments in 3^rd^ instar larva brain of *docrl^KO^*show multiple instances of temporal overlapping [**Fig 6 C,D]**. A representative video of the intracellular calcium imaging for control and *docrl^KO^* is shown in [**Suppl video 4-5].** The mean cross correlation coefficient (CCF) (indicates the absolute lag between two calcium waves of adjacent segments) was lower in *docrl^KO^*than controls at all three temperatures studied [**Fig 6 E]**. We categorised the absolute CCFs as synchronous or asynchronous based on the criteria described in [**Fig 6 B]**. It was observed that at room temperature (22^°^C) *docrl^KO^* have more synchronous pairs (∼67%) compared to controls (∼14%) [**Fig 6F**]. However, the number of calcium waves during the recording duration was not significantly different in *docrl^KO^* compared to control [**Fig 6 G]**.

## Discussion

Human neurodevelopmental disorders are conditions in which structural or functional alterations in normal brain development manifest clinically at birth or in childhood (Raghu et al., 2024). A key challenge in clinically managing neurodevelopmental disorders is the requirement to map the cellular and developmental origins of the condition. One approach is the use of animal models where experimental manipulation can be used to develop and test hypotheses based on observations in human patients. The use of animal models such as rodents can be a powerful approach to model key aspects of the disease phenotype, perform mechanistic analysis and develop potential therapeutics. However, in many cases, rodent models fail to recapitulate the human disease phenotype for a variety of well-understood reasons (Elsea and Lucas, 2002). LS, a neurodevelopmental disorder, exemplifies this scenario where a humanized mouse knockout failed to recapitulate clinical features of the human condition (Festa et al., 2019). In this study, we developed a behavioural assay in a *Drosophila* model of LS that recapitulates febrile seizures, a well described clinical presentation of LS. Germline mutants of *docrl* (*docrl^KO^*) as well as an independent loss of function allele *docrl^crimic^* (Lee et al., 2018) led to heat induced seizures followed by paralysis and a time-dependent recovery. These seizures in *docrl^KO^* could be rescued by a genomic duplication of the X chromosome on the 3^rd^ chromosome whereas in *docrl^crimic^* it could be rescued by expression of *UAS-docrl::gfp.* These findings are similar to that reported by a previous study of heat induced seizures in zebrafish larvae depleted of *OCRL* (Ramirez et al., 2012).

Seizures are a behavioural manifestation of abnormal electrical activity in the brain and can result from both factors intrinsic to brain cells and also as a secondary consequence of metabolic disturbances with primary origin in other organs. In the context of LS, the abnormal renal function and consequent loss of metabolic homeostasis could render the patient susceptible to seizures. Using tissue specific genome engineering, we found that depletion of *docrl* in neurons alone was sufficient to recapitulate heat induced seizures. Conversely, *docrl* depletion in other fly tissues such as nephrocytes and muscles did not lead to this phenotype. It has previously been proposed that altered haemocyte function on OCRL depletion might lead to brain phenotypes (Del Signore et al., 2017).However, our finding that depletion of *docrl* in haemocytes does not lead to heat induced seizures suggests that such a non-cell autonomous signalling from haemocytes to neurons is unlikely to be the mechanism underlying heat induced seizures. Overall, the findings from our study imply that in LS patients, heat-induced seizures arise from a brain intrinsic requirement for the function of this protein in controlling neuronal excitability.

The brain is composed of both glia and neurons and previous studies in flies have implicated both cell types in the origin of heat induced seizures. For example, previous studies have showed that depletion of a voltage-gated potassium channel *sei* (an ortholog of the human ERG channel) in glial cells increases susceptibility to heat-induced seizures (Hill et al., 2019). In contrast, genes such as *cdk8* (a homolog of human *CDK19* that encodes a cyclin dependent kinase) and *sif* (a ortholog of human *TIAM1* that encodes an RAC1 specific guanine exchange factor(GEF)) have been shown to cause bang-sensitive seizures when depleted in neurons but not in glia [Reviewed in (Tanaka and Chung, 2025)]. Our findings in this study clearly show that depletion of *docrl* in glia is insufficient to generate heat induced seizures, whereas depletion in neurons is sufficient to do so. These observations provide compelling evidence that the cell type of origin for seizure susceptibility in *docrl* depleted flies are neurons.

The OCRL protein regulates levels of PI(4,5)P_2_, a lipid that has a well-known role in modulating neuronal excitability. For example, PI(4,5)P_2_ is a key regulator of the synaptic vesicle cycle (Koch and Holt, 2012) and also regulates the activity of multiple ion channels (Hille et al., 2015) and transporters that are known to control neuronal excitability. Thus, the seizures noted on *docrl* depletion could arise from loss of PI(4,5)P_2_ homeostasis in differentiated neurons or from a requirement of PI(4,5)P_2_ in regulating brain development. Using two independent approaches [**Fig 3B and Fig 4C,D**], we found that deletion of *docrl* in neurons, post-differentiation, did not induce these seizures. This finding suggests that the underlying mechanism of the seizures is not solely due to a loss of PI(4,5)P_2_ homeostasis at the plasma membrane of differentiated neurons.

The physiological mechanisms in differentiated neurons that are regulated by *docrl* and underlie heat induced seizure are of interest. Synaptic transmission is a crucial function of neurons and several mutants in synaptic proteins such as *synj*, *nubian*, and *comatose* phenotypically present with seizure-susceptibility (Littleton et al., 1998; Wang et al., 2004). We recorded electroretinograms (A. Nair and P. Raghu, 2011), where the ON/OFF transients that represent synaptic transmission events can be readily monitored. *docrl* depletion did not result in any change in the ON/OFF transients in electroretinograms neither at basal nor elevated temperatures (*data not shown*), implying that this enzyme does not support synaptic phosphoinositide turnover. This is consistent with previous studies that have identified another *Drosophila* lipid phosphatase, *synaptojanin* as the PI(4,5)P_2_ phosphatase that regulates synaptic transmission (Verstreken et al., 2003).

Another critical function of neurons is to form proper functional neural circuits. The larval locomotor circuit has been shown to exhibit a robust delay across ipsilateral segments of the larval brain. One of the mechanisms through which seizures arise is increased synchrony among neuronal populations (Jefferys et al., 2012). This is also observed in larval locomotor circuits, where various seizure mutants show increased synchrony among adjacent ipsilateral segments. Our finding of increased synchrony in *docrl^KO^* at both room temperature and elevated temperatures suggests that defects during early neurogenesis predispose neurons to form improper locomotor circuits that are prone to fire synchronously.

The finding that depletion of *docrl* in differentiated neurons and in adults did not result in seizures suggests that this gene may be required during brain development to prevent heat induced seizures in the adult fly. When might this requirement for *docrl* function be? In *Drosophila*, neurons are generated following the sequential division of neuroblasts and GMC. During the first phase of embryonic neurogenesis, these cell biological events contribute primarily to larval neurons whereas during the second larval phase, following significant remodelling, they generate adult neural circuitry (Chen et al., 2014; Li and Hidalgo, 2020). We found that deletion of *docrl* in neuroblasts and GMC led to adult seizure behaviour and that [**Fig 4 G,H]** temporally controlled depletion of *docrl* during the embryonic 1^st^ phase of neurogenesis was essential to recapitulate the adult seizure phenotype seen in the germline *docrl^KO^*. A previous study using a zebrafish model has presented evidence that depletion of *docrl* during early embryogenesis leads to brain phenotypes including heat induced seizures in fish larvae (Ramirez et al., 2012). Taken together studies in two independent metazoan models find a role for *docrl* in brain development leading to heat induced seizures, a key phenotype in LS patients.

What sub-cellular process might dOCRL regulate during neural stem cell development? Since OCRL is a PI(4,5)P_2_ phosphatase, the levels of both the substrate PI(4,5)P_2_ and the product PI4P are potential links to likely processes. PI(4,5)P_2_ (and OCRL) have been implicated in processes such as vesicular transport including plasma membrane endocytosis [reviewed in (de Sa et al., 2025)], cytoskeletal function (Nández et al.; Vicinanza et al., 2011) and cytokinesis [discussed in (Ben El Kadhi et al., 2012)] while PI4P has been implicated in cell polarity and cytokinesis in multiple model systems (Koe et al., 2018; Xie et al., 2018). In the absence of *docrl*, one or more of these processes, that remain to be discovered, could operate abnormally in neuroblasts and/or ganglion mother cells leading to altered development and abnormal function in adult neurons.

While other 5’ phosphatases, such as s*ynaptojanin* (*synj*) and *dInpp5e*, also exist in *Drosophila* and share an affinity for PI(4,5)P_2_, their functions are distinct. *synj* is highly enriched in synaptic terminals, and its loss leads to defects in synaptic transmission (Vanhauwaert et al., 2017). In contrast, *dInpp5e* is essential for cilia function(Park et al., 2015). Although *synj, docrl*, and *dInpp5e* belong to the same 5’ phosphatase family and all act on PI(4,5)P_2_, they have different developmental requirements and distinct subcellular localizations within the neuron.

Overall, our study establishes the requirement of *docrl* in the brain during early neurogenesis absence of which gives rise to improper neural circuits, which in turn leads to increased susceptibility to seizure in adult flies. In the context of LS patients, it is likely that seizures are a result of a neurodevelopmental defect, and in humans most of the neurogenesis occurs during foetal development (Khodosevich and Sellgren, 2022; Zhou et al., 2023). Thus, potential therapeutic interventions need to be designed bearing this in mind. Our model of LS in *Drosophila* can be used to perform a genetic screen or a drug screening from the LOPAC (Library Of Pharmacologically Active Compounds) database to study the enhancers/suppressors of seizures in *docrl^KO^* mutants.

## Acknowledgements

This study was supported by the Department of Atomic Energy, Government of India (RTI-0046). We thank the Drosophila facility, Imaging Facility and Sequencing facility at NCBS-TIFR for support.

## Materials and Methods

### Fly maintenance

Experiments were performed with *D. melanogaster* strains. All flies were maintained in standard cornmeal media (per litre of media: 80 g corn flour, 20 g D-glucose, 40 g sucrose, 8 g agar,15 g yeast powder, 4 mL propionic acid, 0.6 mL orthophosphoric acid and 1 g methyl 4-hydroxybenzoate dissolved in 5 mL ethanol) at 25^°^C and 70% humidity in a cooled incubator without light-dark cycles. For behavioural experiments, flies were grown in jaggery-rich media (per litre of media: 80 g corn flour, 70 g jaggery, 9 g agar, 15 g yeast powder, 4.4 mL propionic acid and 1.25 g methyl 4-hydroxybenzoate dissolved in 5 mL ethanol), and for experiments with larvae, control and germline *docrl^KO^*were grown on jaggery-rich media supplemented with yeast paste. A full list of fly strains used in this study is given in Supplementary Table 1.

### Behaviour Assays

#### Seizure Behaviour

Adult flies were collected at eclosion in a fresh vial and aged for 2-4 days. On the day of the experiment, 5 flies (sex sorted) were transferred into an empty vial and allowed to recover from carbon dioxide (CO_2_) anaesthesia for 1 hour. Vials containing flies are lowered in water at 42^°^C in a 2 L beaker for *docrl^KO^* and *docrl^crimic^* heat shock was given for 45 sec-1min and for tissue-specific knockouts a 3 min heat shock was used. The heat shock protocol was adapted from (Hill et al., 2019; Mituzaite et al., 2021). Movies of fly behaviour were collected using a Logitech c615 full HD webcamera along with OBS studio at 30fps or by phone camera for better resolution, video files were stored in mkv format and used for subsequent analysis. Each experiment was composed of 5 flies/vial and two trials were conducted per set. Analysis was done using VLC media player by manually counting the flies in the video file and recording them in Excel sheets. These Excel sheets contain time and the fraction of flies ranging from 0-1 for the parameter measured during seizure induction-called falling fraction. After seizure induction, the vial is taken out from the water bath (within 10s) and counted for the number of flies paralysed-called paralysis fraction. Vials are observed at defined time intervals for the number of flies recovered from paralysis-called recovered fraction. Definition of falling frequency: The number of flies (seizing or fallen) at the base of the vial/ total number of flies in the vial during seizure induction calculated every 10s of the movie. Paralysis fraction: Number of flies that are unable to maintain their upright posture after seizure induction/by total number of flies at time 1 min for *docrl^KO^, docrl^crimic^* and 3 min for tissue specific knockouts in the video. Recovery fraction: The fraction of flies moving after paralysis, counted every 10s for 6-10 mins after the vial was taken out of the water bath.

### Physiological assays

#### Calcium imaging

Brains of 3^rd^ instar late feeding larvae were dissected in physiological saline (PS) (Pulver et al., 2015; Streit et al., 2016), embedded on a coverslip with 0.5-1 % low melting agarose and 10 ml of PS was added on top. Fictitious calcium waves originating in the motor neuron ganglia were imaged using a 20X objective on an Olympus IX71 Microscope (RRID:SCR_022185) equipped with a GFP filter in LUDL MAC6000 controlled by µmanager (Edelstein et al., 2010). An XCITE 120LED light source was used for excitation and images were collected on a QImaging Retiga 6000 camera. Temperature control of the microscope stage was provided by the Tokai Hit thermo plate. Images were taken at 5 fps for 600 frames (∼2mins). From the image files, ROIs (Region of interest) were marked using FIJI and intensity profiles across time were obtained. The intensity profiles for each ROI were analysed using a custom-written Python code. Intensities were normalised to the minima of fluorescence intensities measured across time. The cross correlation across two adjacent ipsilateral ROI was determined using xcorr function in matplotlib library and the number of firings were calculated by the find peaks function in scipy library. To generate a category plot, synchrony was considered if the cross-correlation value lies between −/+200ms.

### Molecular Biology

#### Polymerase chain reaction (PCR)

Flies were homogenized in 40ml of squishing buffer (10 mM Tris-HCl pH 8, 1 mM EDTA, 25 mM NaCl, 200 g/ml Proteinase K) or fly heads were squished in 3ml/head squishing buffer. Lysates were incubated at 37^°^C for 30 min and then at 95^°^C 3 min. 2ml of lysate was used to perform PCR with respective primer sets. Primers used are as follows

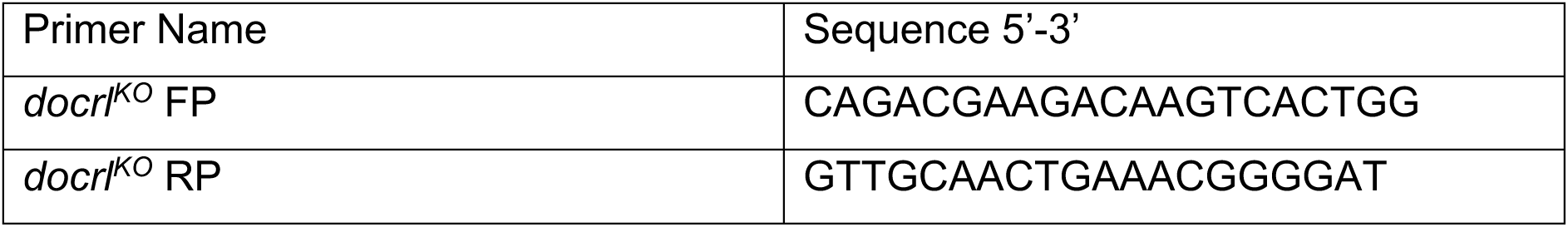

#### Western Blotting

20 adult brains were dissected into 15ml of ice cold larval lysis buffer (50 mM Tris-HCl pH 7.5, 1 mM EGTA, 1 mM EDTA, 1% Triton X-100, 50 mM NaF, 0.27 M sucrose, 0.1% β-Mercaptoethanol) and homogenised in a bath sonicator for 1 min or a whole fly was homogenised in 30mL of 2X Laemmli loading buffer for whole fly lysate Homogenized samples were heated with 2X Laemmli loading buffer for 5 min at 95^°^C. Samples were separated using a 10% acrylamide gel using standard procedures. Following electrophoresis, proteins were electro-transferred onto a 0.45µm nitrocellulose membrane. Membranes were blocked using 5% Blotto in PBS+0.1% Tween-20 (PBST). Blocked membranes were incubated with a primary antiserum against dOCRL (Del Signore et al., 2017)(a gift from Avital Rodal) (1:1500 dilution in 5 % Blotto in PBST) overnight at 4^°^C. As a loading control β-tubulin was detected using DSHB Cat# E7, RRID:AB_528499 (1: 4000 dilution in 5% Blotto in PBST). Following washes in PBST membranes were detected using horseradish peroxidase conjugated secondary antibody against respective primary antibody ( Jackson Laboratories, Pennsylvania, USA) used in 1:10000 dilution for 2 hr at room temperature. Following washes in PBST, the membrane was imaged using an Invitrogen iBright FL1500 Imaging System (RRID:SCR_026331) or GE Healthcare ImageQuant™ LAS 4000 following incubation with an enhanced chemiluminescence substrate (BioRad Cat #170-5061).

### Immunohistochemistry and Imaging

For embryo staining, eggs were laid for 4 hrs and aged for 12-14 hrs to achieve stage 16-17. Staining was adapted from (https://denisemontell.mcdb.ucsb.edu/sites/default/files/2021-10/collecting-and-staining-embryos.pdf). Embryos were dechorionated by 4% bleach (Qualigens #CAS 7681-52-9) for 2-3 min, followed by vigorous shaking for 30s in 1:1 4% Paraformaldehyde (PFA)(Electron Microscopy Sciences Cat#15710) and N-heptane. Embryos were left for 25 mins for fixation. Equal amount of methanol was added to the microcentrifuge tube and devitellinization was done by gentle shaking. Embryos sunk to the bottom of the microcentrifuge tube were washed twice with methanol, rehydrated by washing with PBS+0.3% TritonX-100 (PBTX) thrice and blocked with 10% NGS (normal goat serum, HIMEDIA Cat#RM10701) in PBTX (blocking solution) for 1hr at RT. Embryos were incubated with respective primary antibodies [chicken α-GFP (1:4000) (Abcam Cat# ab13970, RRID:AB_300798); rat α-Elav (1:200)(DSHB Cat#Rat-Elav-7E8A10, RRID:AB_528218)] overnight at 4^°^C followed by four washes with PBTX for 10 mins each. Embryos were incubated with Alexa Fluor™ conjugated secondary antibody [goat α-chicken Alexa Fluor™ 488 (1:500)( Thermo Fisher Scientific Cat# A-11039, RRID:AB_2534096); goat α-rat Alexa Fluor™ 568 (1:500)( Thermo Fisher Scientific Cat# A-11077, RRID:AB_2534121)] for 4hrs at RT. Followed by four washes with PBTX for 10mins each. Embryos were mounted on a glass slide with spacers made by two pieces of double sided tape in 90% glycerol and covered with a No.1 coverslip. Image acquisition was done in Olympus Confocal Laser Scanning Microscope Fluoview FV3000 (RRID:SCR_017015) with 40X objective (NA=1.3).Images were analysed using Fiji (Schindelin et al., 2012).

For wandering 3^rd^ instar and adults, brains were dissected in ice cold PBS and fixed with 4% PFA for 25 mins at RT. Brains were permeabilised by washing in PBTX (twice). Blocking solution was added and brief freezing in −20^°^C for 20 min was done to improve permeabilisation (Thapa et al., 2019) followed by 40 mins at RT. Brains were incubated in primary antibodies [chicken α-GFP(1:4000)( Abcam Cat# ab13970, RRID:AB_300798), rat α-Elav (1:200)( DSHB Cat#Rat-Elav-7E8A10, RRID:AB_528218),mouse α-Elav(1:100)( DSHB Cat# Elav-9F8A9, RRID:AB_528217), rat α-Deadpan(1:100)( Abcam Cat# ab195173, RRID:AB_2687586)] in blocking solution for 24 hrs at 4^°^C followed by four washes with PBTX for 10mins each. Then the brains were incubated with respective Alexa Fluor™ conjugated secondary antibodies [goat anti-chicken Alexa Fluor™ 488 (1:500)( Thermo Fisher Scientific Cat# A-11039, RRID:AB_2534096);goat anti-rat Alexa Fluor™ 568(1:500)( Thermo Fisher Scientific Cat# A-11077, RRID:AB_2534121); goat anti-mouse Alexa Fluor 633(1:500)( Thermo Fisher Scientific Cat# A-21052, RRID:AB_2535719)] for 4 hrs at RT. Following four washes with PBTX for 10mins each, brains were mounted on a glass slide with spacers made by two pieces of double sided tape in 90% glycerol and covered with a No.1 coverslip. Image acquisition was done in Olympus Confocal Laser Scanning Microscope Fluoview FV3000 (RRID:SCR_017015) with 40X objective (NA=1.3) or 20X objective (NA=0.85). Images were analyzed using Fiji (Schindelin et al., 2012).

### Statistical analysis

For seizure behaviour analysis, statistics were only used for paralysis fraction. Most of the time, normality tests like the Sapiro-Wilk test and Q-Q plot analysis failed in our behavioural data; hence to obtain a consistent statistical significance we used a non-parametric tests like the Mann-Whitney U test (to compare across two groups) and the Kruskal-Wallis test, post hoc Dunn’s test for multiple comparison (to compare across more than two groups). A similar strategy was used for intracellular calcium imaging and confocal data. All the graphs were generated and statistics performed in GraphPad Prism version 10.2.0, GraphPad Software, Boston, Massachusetts USA.

## Supplementary figure legends

**Supp Figure 1: (A)** PCR analysis for validation of tissue specific knockout of *docrl* by CRISPR/Cas9. The genotypes are germline *docrl^KO^* (*docrl^KO^/FM7i,gfp*) and ubiquitous somatic cell *docrl* knockout by expression of *da-GAL4 i.e. docrl^K-D^ (dgRNAs;UAS-Cas9-T2A-egfp/da-GAL4)* and the negative control i.e. *da>* (*da-GAL4).* Arrowhead indicates the diagnostic band detecting the *docrl* genomic deletion. **(B)**Western blot detecting the depletion of dOCRL across most larval cells using CRISPR/Cas9. The genotypes shown are positive control *da>* (*da-GAL4),* ubiquitous somatic knockout *docrl^K-D^ (dgRNAs;UAS-Cas9-T2A-egfp/da-GAL4).* Germline *docrl^KO^* is used to validate the dOCRL protein band detected by the polyclonal antibody we have used.**(C-E)** Heat induced seizure assay 2-4 days old adult flies. **(C)** The X axis represent time in seconds. The Y axis represents the falling fraction **(D)** paralysis fraction and **(E)** recovery fraction respectively. The genotypes are control (*da-GAL4*) (N=23 sets, n=115 flies), *docrl^K-D^(dgRNAs;UAS-Cas9-T2A-egfp/da-GAL4) (*N=19 sets,n=95 flies). The graph shows mean±95% CI, Statistics used : Mann-Whitney U test,**** represents p value <0.0001.

**Suppl Figure 2:** Heat induced seizure assay 2-4 days old adult flies in which *docrl* has been depleted in specific tissues. The X axis represents the time in seconds The Y axis represents the falling fraction. Error bars show mean falling fraction ±95% CI. The genotypes of control (shown in black color) (xx*-GAL4;dgRNAs*) (N=6-10 sets, n=30-50 flies) where xx is the tissue specific promoter, *docrl* knockout in tissue(shown in pink color*) (xx-GAL4;dgRNAs;UAS-Cas9-T2A-egfp) (*N=6-10 sets, n=30-50 flies).The knockout in **(A)** nephrocyte using *dot-GAL4 (docrl^K-R^);* **(B)** eye using *GMR-GAL4 (docrl^K-E^);* **(C)** salivary gland using *AB1-GAL4 (docrl^K-SG^);* **(D)** haemocyte using *hml-GAL4 (docrl^K-H^);* **(E)** fat body using *lpp-GAL4 (docrl^K-FB^);* **(F)** muscle using *Mef2-GAL4 (docrl^K-M^)*.

**Suppl Figure 3:** Heat induced seizure assay 2-4 days old adult flies in which *docrl* has been depleted in specific subsets of neurons. The X axis represents the time in seconds The Y axis represents the falling fraction. Error bars show mean falling frequency±95% CI. The genotypes of control (shown in black color)(xx*-GAL4;dgRNAs*) (N=5-12 sets, n=25-60 flies) where xx is the neuronal specific promoter, *docrl* knockout in neuronal subtype (shown in pink color) *(xx-GAL4;dgRNAs;UAS-Cas9-T2A-egfp) (*N=5-13 sets, n=25-65 flies). The neuronal subtypes are **(A)** Glutamatergic **(B)** Serotonergic and Dopaminergic **(C)** Cholinergic **(D)** GABergic **(E)** Octopaminergic **(F)** Peptidergic. The GAL4 lines used for each subclass are described in materials and methods.

## Supplementary Videos

**Suppl Video 1:** A representative video of heat induced seizure assay with germline *docrl^KO^* adult flies. The temperature of the water bath is 42^°^C. The germline *docrl^KO^*(*docrl^KO^/Y)* flies are shown in pink, the controls (*y^1^v^1^/Y)* and the rescue (*docrl^KO^/Y;;Dp(1:3)DC402*) are shown in green. A timer below shows the time elapsed in the water bath. The yellow text on the right hand side shows the parameters quantified from the assay. During paralysis the *docrl^KO^* flies are not moving and lose their posture compared to controls.

**Suppl Video 2:** A representative video of heat induced seizure assay with pan neuronal *docrl* knockout adult flies. The temperature of the water bath is 42^°^C. The genotypes of the control (*elav-GAL4;dgRNAs)* flies are shown in black, the pan neuronal knockout *docrl^K-N^* (*elav-GAL4;dgRNAs;UAS-Cas9-T2A-egfp)* in pink. The timer below shows the time elapsed in the water bath. The yellow text on the right hand side shows the parameters quantified from the assay.

**Suppl Video 3:** A representative video of heat induced seizure assay with pan glial *docrl* knockout adult flies. The temperature of the water bath is 42^°^C. The genotypes of the control (*dgRNAs;repo-GAL4)* flies are shown in black, the pan glial knockout *docrl^K-G^* (*dgRNAs;UAS-Cas9-T2A-egfp/repo-GAL4)* in pink. The timer below shows the time elapsed in the water bath. The yellow text on the right hand side shows the parameters quantified from the assay.

**Suppl Video 4:** A representative video showing intracellular calcium imaging of 3^rd^ instar larval brain in controls (*y^1^v^1^;ok6-GAL4;UAS-GCaMP5G*); expressing the calcium sensor GCaMP5G under a motor neuron specific GAL4 (*ok6-GAL4).* Two circular, adjacent ipsilateral ROIs (ROI), marked in blue and red, are shown in the video. Their respective normalised intensity profiles are shown on the right, with the X-axis representing time and the Y-axis representing the normalised intensity values across a 50-frame running window.

**Suppl Video 5:** A representative video showing intracellular calcium imaging of 3^rd^ instar larval brain in *docrl^KO^* (*docrl^KO^;ok6-GAL4;UAS-GCaMP5G*); expressing calcium sensor GCaMP5G under a motor neuron specific GAL4 (*ok6-GAL4).* Two circular, adjacent ipsilateral ROIs (ROI), marked in blue and red, are shown in the video. Their respective normalised intensity profiles are shown on the right, with the X-axis representing time and the Y-axis representing the normalised intensity values across a 50-frame running window.

**Supplementary Table S1.**

| Strain | Description | Source | Reference |
| --- | --- | --- | --- |
| Oregon -R | Wild type | RRID:BDSC_2376 |  |
| *y<sup>1</sup>v<sup>1</sup>* | Genetic control for *docrl<sup>KO</sup>* | In Lab |  |
| *docrl<sup>KO</sup>* | Germline *docrl<sup>KO</sup>* | RRID:BDSC_93740 | (Trivedi et al., 2020) |
| *Dp(1;3)DC402* | Duplication of X chromosome on 3<sup>rd</sup> | RRID:BDSC_31454 | (Del Signore et al., 2017) |
| *docrl<sup>crimic</sup>* | CRIMIC fly of *docrl* | RRID:BDSC_83167 | (Lee et al., 2018) |
| *UAS-docrl::gfp* | gfp tagged *docrl* | Avital Rodal's Lab | (Del Signore et al., 2017) |
| *dgRNAs* | Dual guide RNAs for *docrl* | RRID:BDSC_92492 | (Trivedi et al., 2020) |
| *UAS-Cas9-T2A-egfp* | GAL4 dependent Cas9 expression with a self-cleavable egfp | RRID:BDSC_94298 | (Trivedi et al., 2020) |
| *elav-GAL4* | GAL4 expressing in post mitotic neurons | RRID:BDSC_458 | (Hill et al., 2019) |
| *repo-GAL4* | GAL4 expressing in glia | RRID:BDSC_7415 | (Hill et al., 2019) |
| *GMR-GAL4* | GAL4 expressing in photoreceptors | RRID:BDSC_1104 | (Basak et al., 2021) |
| *dot-GAL4* | GAL4 expressing in nephrocytes | RRID:BDSC_6903 | (Ramesh et al., 2024) |
| <u>*AB1-GAL4*</u> | GAL4 expressing in salivary glands | RRID:BDSC_1824 | (Sharma et al., 2019) |
| *lpp-GAL4* | GAL4 expressing in fatbody | RRID:BDSC_84317 | (Zappia et al., 2021) |
| *mef2-GAL4* | GAL4 expressing in muscles | RRID:BDSC_27390 | (Zappia et al., 2021) |
| *hml-GAL4* | GAL4 expressing in haemocytes | Tina Mukherjee Lab | (Sinenko and Mathey-Prevot, 2004) |
| *nSyb-GAL4* | GAL4 expressing in mature neurons | BDSC_51635 | (Kobler et al., 2021) |
| *grh-GAL4* | GAL4 expressing in neuroblast | Gaiti Hasan Lab | (Almeida and Bray, 2005) |
| *pros-GAL4* | GAL4 expressing in GMCs | Gaiti Hasan Lab | (Ohshiro et al., 2000) |
| *elav-GAL4;;tub-GAL80<sup>ts</sup>* | GAL4/GAL80<sup>ts</sup> for neurons | Flyfacility, NCBS, Bangalore |  |
| *elav<sup>GS</sup>* | Geneswitch GAL4 expressing in neurons | RRID:BDSC_43642 | (Osterwalder et al., 2001) |
| *ok6-GAL4* | GAL4 expressing in motor neurons | RRID:BDSC_64199 | (Sanyal, 2009) |
| *UAS-GCaMP5G* | Genetically encoded calcium sensor | RRID:BDSC_56500 | (Akerboom et al., 2012) |
| *da-GAL4* | GAL4 expressing in most of the cells | In lab | (Gupta et al., 2013) |
| *vglut-GAL4* | GAL4 expressing in glutamatergic neurons | RRID:BDSC_24635 | (Aimon et al., 2023) |
| *ddc-GAL4* | GAL4 expressing in serotonergic and dopaminergic neurons | RRID:BDSC_7010 | (Reiszadeh Jahromi et al., 2021) |
| *chat-GAL4* | GAL4 expressing in cholinergic neurons | RRID:BDSC_56500 | (Hussain et al., 2024) |
| *gad1-GAL4* | GAL4 expressing in GABAergic neurons | RRID:BDSC_51630 | (Hill et al., 2019) |
| *tbh-GAL4* | GAL4 expressing in octopaminergic neurons | RRID:BDSC_39939 | (Hill et al., 2019) |
| *c929-GAL4* | GAL4 expressing in peptidergic neurons | RRID:BDSC_25373 | (Hill et al., 2019) |

