## Supplementary figures and images for "Neurodevelopmental origin of seizures in Lowe syndrome"

### Supplementary Figures

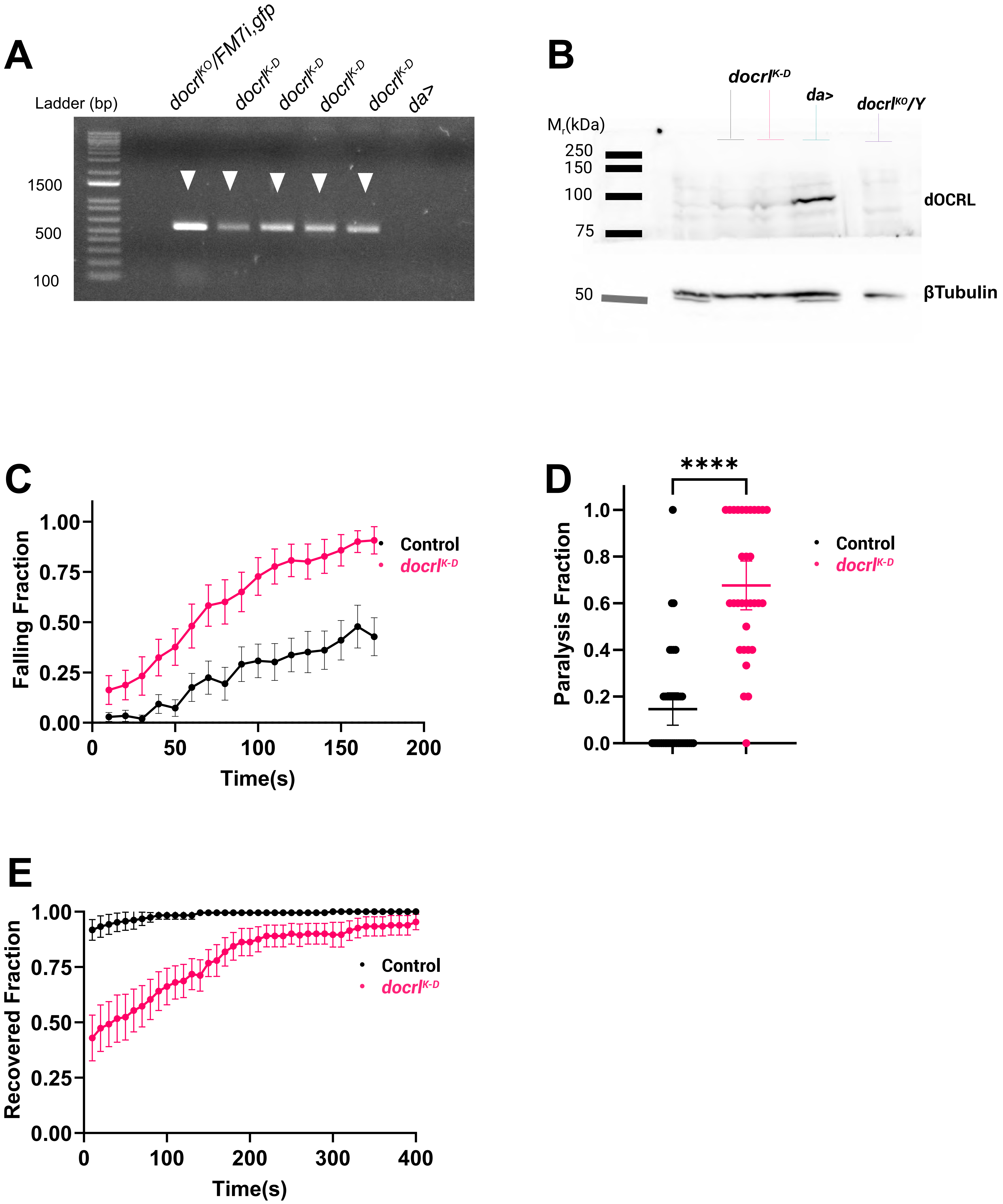


Behera et.al Suppl Figure 1

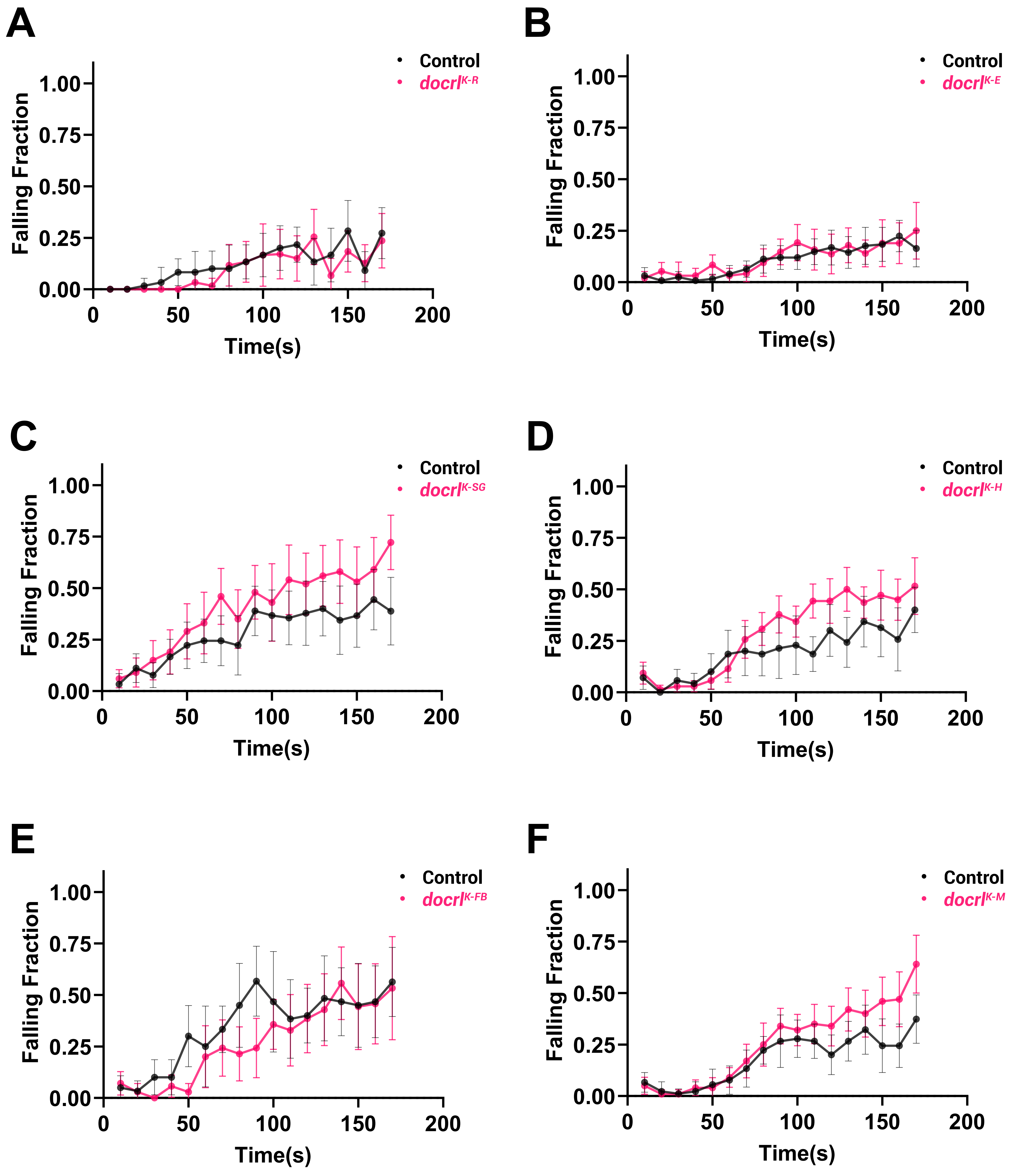

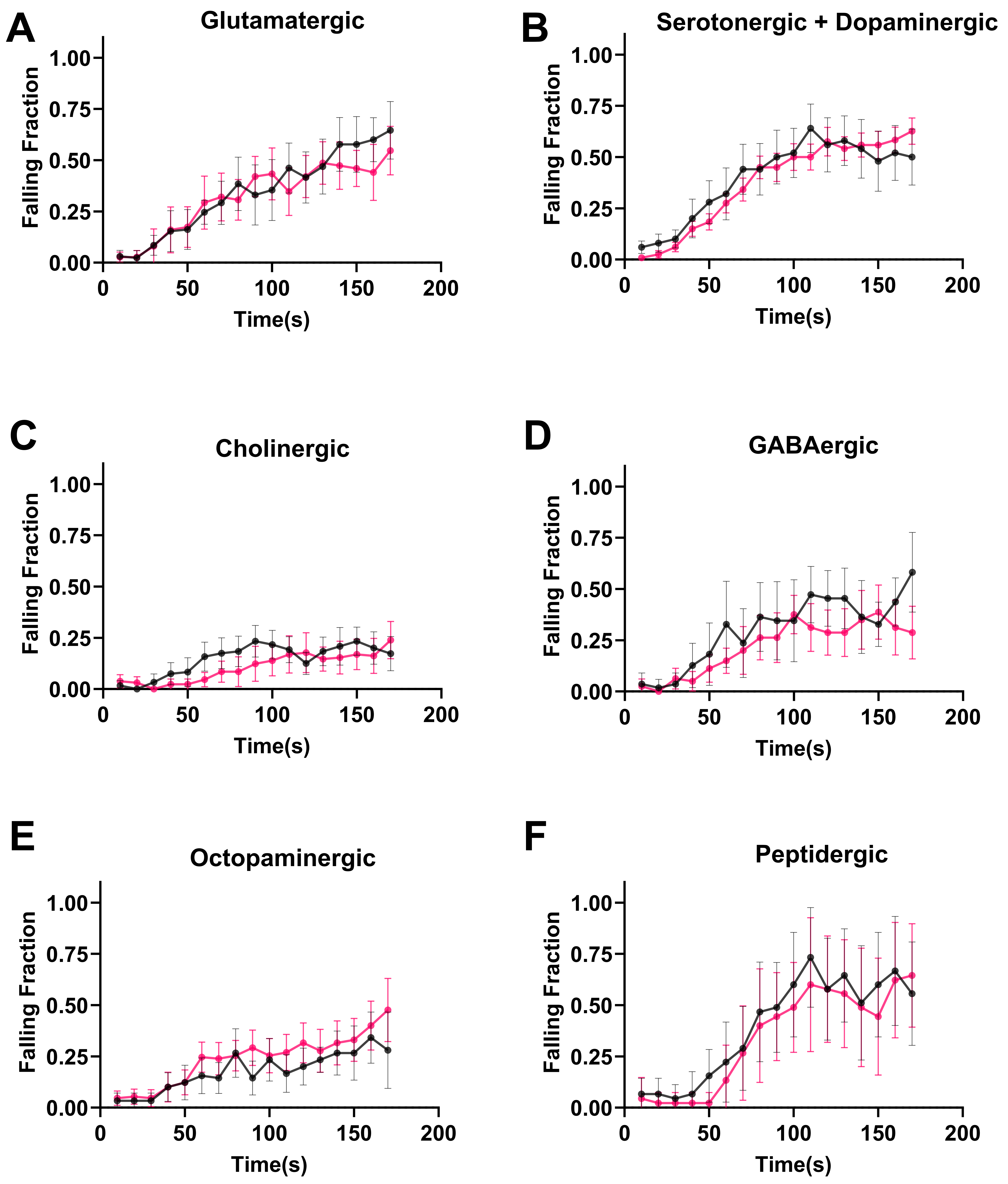


Behera et.al Suppl Figure 3
